# THE EFFECT OF SERTRALINE AND VOLUNTARY EXERCISE DURING PREGNANCY ON LITTER CHARACTERISTICS AND POSTNATAL AFFECTIVE BEHAVIOUR IN RAT DAMS

**DOI:** 10.1101/2025.02.02.636079

**Authors:** Noor S. Jarbou, Olivia Mairinger, Elise Kulen, Lucas Mushahwar, Samara Walpole, Jasmine Matthews, Simon Maksour, Katrina Weston-Green, Mirella Dottori, Kelly A. Newell

## Abstract

**Introduction:** Sertraline is the frontline pharmacotherapy for the treatment of depression and anxiety during pregnancy. However, there is little evidence regarding the effects of sertraline on maternal behaviour or the maternal brain. Furthermore, the efficacy of non-pharmacological approaches to treatment in pregnancy, such as exercise, are unclear. Therefore, the aim of this study was to examine the effects of sertraline and exercise during pregnancy on maternal postnatal depressive-like, anxiety-like and associated behaviours, as well as litter characteristics, in a rat model of depression. We also investigated the effects of these treatments on the maternal brain, focusing initially on DNA methylation and glutamatergic markers, which have been implicated in depression.

**Method:** Twenty-four female Wistar-Kyoto (WKY; strain that models depression and anxiety) rats were divided into three groups: 1. WKY-Sertraline; 2. WKY-Exercise, 3. WKY-Vehicle; Six female Wistar rats were included as controls. Rats were treated with sertraline (10mg/kg) or vehicle (33% propylene glycol) twice/day, from gestational day (GD) 1 to postnatal day (PN) 14. The WKY-Exercise group were provided access to a running wheel during pregnancy for 3 hours/day from GD1-18. Dam and litter characteristics, as well as pup ultrasonic vocalisations (USVs), were measured. Dams underwent behavioural testing at 5-weeks postnatal to assess depressive-, anxiety- and cognitive-like behaviours. Gene expression of DNA methylation markers (*Dnmt1, Dnmt3a*) and glutamate receptors (*Grin1, Grin2a, Grin2b*) were measured in the prefrontal cortex (PFC), using RT-qPCR.

**Result:** The WKY-Sertraline group gained 39% less weight in their first pregnancy week compared to all other groups (*p*<0.05) and produced smaller litters compared to Wistar controls (-43%; *p*=0.003) and WKY-Exercise (-38%; *p*=0.012), and WKY-Sertraline pups had slightly smaller brain weights (*p*=0.031 compared to WKY-Vehicle). The WKY-Exercise pups produced increased number of USVs at PN7 compared to WKY-Vehicle, with no treatment differences at PN14. The WKY strain however, did showed reduced average mean amplitude of calls and reduced average duration of calls at PN7 compared to WIS (p<0.01) and reduced number of calls at PN14 (p<0.01). Maternal sertraline treatment did not significantly affect dam behavioural measures, all maternal cortical gene expression. The WKY-Exercise group however showed reduced anxiety-like behaviours, spending more time in the open arms (620%; *p*=0.027) and less time in the closed arms (-22%; *p*=0.047) of the elevated plus maze (EPM) compared to WKY-Vehicle, and more time in the centre of the open field test (OFT) compared to WKY-Vehicle (132%; *p*=0.057). Furthermore, WKY-Exercise dams showed a 64% increase in *Dnmt3a* mRNA levels in the PFC compared to WKY-Vehicle (p=0.019).

**Conclusion:** Voluntary exercise during pregnancy in the WKY rat model, reduced postnatal anxiety-like behaviour in the dam. This was accompanied by elevated *DNMT3a* gene expression in the PFC, suggesting this region may be sensitive to DNA methylation changes following maternal exercise. In contrast, maternal sertraline did not impact these behaviours or genes. Maintaining sertraline treatment beyond PN14, may have resulted in broader effects on dam behaviour, which should be explored further. Maternal sertraline did appear to have some adverse effects on the *in utero* environment, evidenced by smaller litters, with slightly smaller pup brain weights, which should be investigated further. Our findings suggest a long-term beneficial effect of exercise during pregnancy and supports future studies examining the effects of exercise in antenatal depression in the human population.

## 1. Introduction

Depression and anxiety during pregnancy have a worldwide prevalence of approximately 20% of the pregnant population (Albertini et al., 2024; Amani et al., 2021; Dennis et al., 2017; Fawcett et al., 2019). Untreated or unmanaged antenatal depression or anxiety can have serious negative outcomes for both mothers and babies, such as poor self-care, premature labour, low birth weight, longer hospital stays and weakened mother-child bonds (Dunkel Schetter and Tanner, 2012; Field, 2011; Li et al., 2021). It has been estimated that between 3-10% of pregnant women are prescribed antidepressant drugs (Cooper et al., 2007; Huybrechts et al., 2013; Molenaar et al., 2020), most frequently selective serotonin re-uptake inhibitors (SSRIs) (ACOG, 2023; Molenaar et al., 2018). While there are many SSRIs available, clinical practice guidelines in Australia, New Zealand and the United States of America recommend that following psychotherapy, the SSRI sertraline is prescribed as the first-line treatment for antenatal depression and anxiety, where there is no history of successful pharmacotherapy prior to pregnancy (ACOG, 2023; Molenaar et al., 2018).

SSRIs, including sertraline, act by blocking the serotonin transporter and inhibiting the re-uptake of serotonin into the presynaptic neuron, resulting in increased serotonin in the synaptic cleft (Hiemke and Härtter, 2000). SSRIs and their metabolites cross the placenta and can be secreted in breast milk, exposing the child before and after birth (Heikkine et al., 2002; Rampono et al., 2004). Despite the relatively high rate of women that are prescribed antidepressant drugs during their pregnancies, to our knowledge, there have been limited studies assessing the effectiveness of treatment with SSRIs (or other antidepressant medications) during pregnancy on the mother’s behaviour. Additionally, we know very little about how these medications affect the maternal brain. Pioneering work from the Pawluski lab showed that postnatal treatment (PN1-PN28) with the SSRI, fluoxetine, in a model of gestational stress, prevented the stress-induced increase in anxiety-like behaviour in dams, but had no effect on immobility time in the forced swim test (Pawluski et al., 2012). Fluoxetine treatment following gestational stress was also shown to increase hippocampal neurogenesis in dams (Pawluski et al., 2012). Gemmell et al. (2016) extended these findings to show that fluoxetine treatment during gestation and postnatally (GD15-PN21) did not alter maternal caregiving behaviours (Gemmel et al., 2016), but reduced the gestational stress-induced increases in cortical synaptophysin density and decreased the density of the epigenetic marker, DNA methyl transferase 3a (DNMT3a), in the dentate gyrus of the maternal hippocampus (Gemmel et al., 2016). However, it is not clear if the SSRI sertraline has similar behavioural or neurobiological effects when used during pregnancy.

There is growing evidence that exercise has the ability to alleviate symptoms of clinical depression and anxiety in the general (non-pregnant) population (Belvederi Murri et al., 2018; Kandola and Stubbs, 2020). It is well known that exercise during pregnancy has beneficial effects on the general health and wellbeing of the mother and her infant (Kołomańska et al., 2019). However, research into the effects of exercise in pregnancy for the treatment of maternal anxiety and depression is limited (for review see Jarbou and Newell, 2022). One rodent study, using a corticosterone-induced postpartum depression rat model, showed that voluntary running wheel exercise over eight weeks (including, pre-gestation, gestation, postnatal) decreased depressive-like behaviour in the forced swim test on postnatal day 22 and 23 and increased markers of hippocampal neurogenesis (Gobinath et al., 2018). However, whether exercising during pregnancy specifically can alter postnatal dam behaviour or the postnatal brain has not been examined.

This study therefore aimed to determine the effects of sertraline and exercise during pregnancy on dam postnatal affective behaviours and cortical and hippocampal gene expression, in a rodent model relevant to depression and anxiety. Furthermore, we aimed to determine how these treatments affect litter and pup characteristics.

## 2. Methodology

### 2.1 Animal housing

All experiments were approved by the University of Wollongong Animal Ethics Committee and were carried out in accordance with the Australian Code for the Care and Use of Animals for Scientific Purposes (AE20/11). Twenty-four female and 10 male Wistar-Kyoto (WKY) rats and 6 female and 2 male Wistar (Wis) rats were obtained from Animal Resources Centre (ARC) in Perth, Western Australia. The WKY rats are an inbred rat strain that demonstrates a depressive-like and anxiety-like phenotype as per our previous studies (Millard et al., 2021, 2020, 2019) and the WIS rats are widely used as a control strain (Rao and Sadananda, 2015; van Zyl et al., 2014; Wang et al., 2023). All rats were housed at the University of Wollongong animal housing facility. Rats were housed 2/cage of the same sex and strain in Techniplast GR1800 Double Decker individually ventilated cages (IVC) with corncob bedding. The rats were housed in a temperature-controlled environment (20±2C°) under reverse lighting (lights off at 7:00 am) to capture behavioural measures during the rat’s nocturnal activity. Rats had free access to lab chow and water.

Rats were allowed at least one week to habituate to their new environment followed by one week of training the female rats to eat 1/8^th^ of a wafer biscuit (Evropa Jadran Wafer, Macedonia). All female rats underwent baseline behavioural testing including the open field test (OFT), novel object recognition (NOR) and elevated plus maze (EPM) before mating. During the mating period, a single female rat was housed with a single male of the same strain for four days, resulting in a 100% mating success rate as per our previous study (Millard et al., 2019). After the mating period (four days), females were returned to their home cage with their original female cage mate of the same strain and group. Female WKY rats were randomly distributed into three groups, 8 rats/group: Vehicle (WKY-Veh), Sertraline (WKY-Sert), Exercise (WKY-Ex), and all Wis rats were assigned to vehicle group only. At gestational day 19-21, females were housed 1/cage in preparation for birth. Following the birth of pups, which was designated as postnatal day 0 (PN0), pups were housed with their dam and littermates until approx. PN28, at which time they were weaned and housed 2-3 per cage with their littermates of the same sex. The dams were then re-housed 2/cage with their original cage mate of the same group.

### 2.2. Animal treatment and exercise protocol

The pregnant WKY-Sert rats were treated twice a day with sertraline hydrochloride (S0507, Tokyo Chemical Industry Co.) at a dose of 10 mg/kg (total of 20 mg/kg/day), as per previous studies (Kott et al., 2019; Lozano et al., 2021; Pawluski et al., 2020). Sertraline was dissolved in vehicle (66% propylene glycol), injected in 1/8^th^ of a wafer biscuit, and offered to the rats twice a day, due to the elimination half-life of sertraline hydrochloride being approximately 5 hours in rats (Chouinard, 1992; Pereira-Figueiredo et al., 2014). Pregnant rats in the WKY-Veh, WKY-Ex and Wis groups were similarly treated with the wafer biscuits containing only vehicle. Rats were monitored to ensure the wafer biscuit was consumed. The selected administration method was chosen as it models the clinically relevant scenario in which sertraline is orally administered, it minimises stress to the rats which can result from injection, and is accepted in the literature (Pawluski et al., 2020; Singh and Saadabadi, 2021). The treatment regimen commenced at the end of the mating period, continued throughout the gestational term and was discontinued at PN14 as per previous studies in our group (Millard et al., 2021, 2019) (Figure 1).

**Figure 1:**
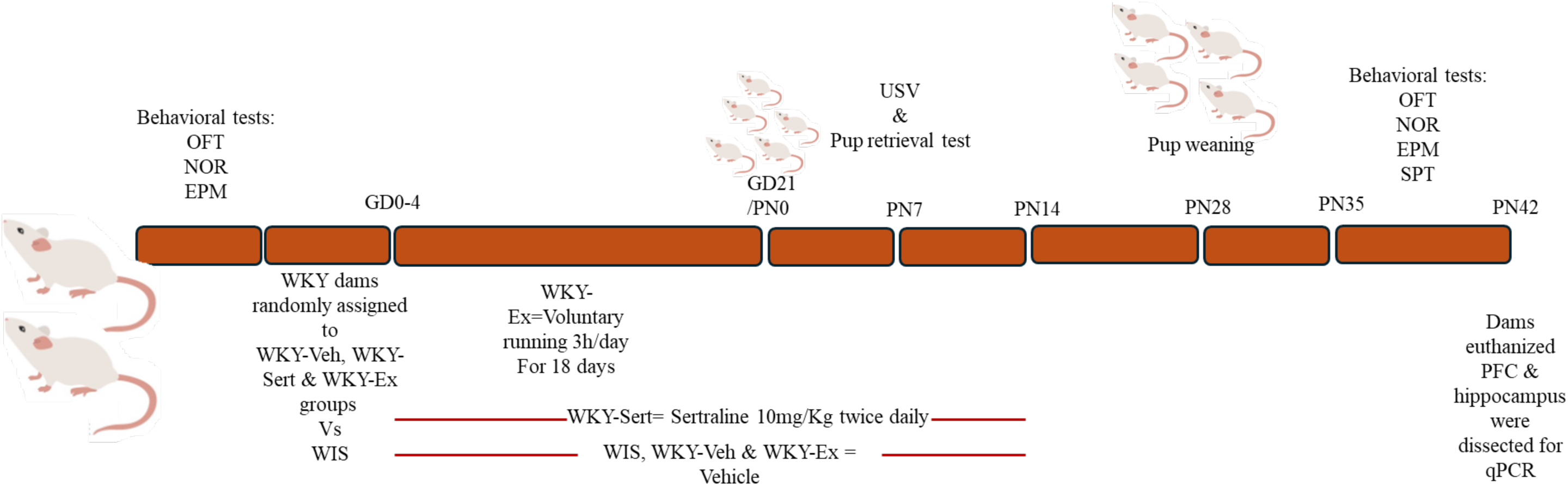
Study timeline and treatment paradigm. Mating period=4 days. Ex: exercise; GD: gastational day; EPM: elevated plus maze; NOR: novel object recognition; OFT: open field test; PFC: prefrontal cortex; PN: postnatal day; Sert: sertraline; SPT: sucrose preference test; USV: ultrasonic vocalizations; Veh: vehicle; Wis: Wistar; WKY: Wistar-Kyoto.

After the mating period, rats in the WKY-Ex group were placed 1 dam/cage in cages equipped with a freely spinning running wheel connected to an integrated living chamber (Activity Wheel and Living Chamber, Model 80859, Lafayette Instrument Company, IN, USA) (Babic et al., 2018), for 3 hours/day from 11:00 am-2:00 pm for 18 days, to allow voluntary running. Each cage contained corncob bedding and provided free access to food and water. An infrared sensor counter was attached to a USB computer interface to record the distance travelled (in metres), then it was quantified using Activity Wheel Monitor Software (Lafayette Instrument Company, IN, USA).

### 2.3. Litter characteristics and behavioural tests

Dam body weight was recorded daily during the treatment period, from the end of the mating period to PN14. Litter size and pup body weights were recorded at PN7 and PN 14 and the sex of the pups was confirmed on PN14. The pup retrieval test was performed at PN7, to assess mum-pup interactions (maternal behaviour). Ultrasonic vocalization (USV) recordings were conducted on the pups at both PN7 and PN14. 1-2 pups/sex were euthanized from each dam at PN14, by rapid decapitation. The pup brains were dissected and weighed. At PN35, after pups were weaned and dams were housed with their original cage mate, dams underwent a series of behavioural tests including the OFT, NOR, EPM and sucrose preference test. The OFT, NOR and EPM apparatus were cleaned between each rat using 70% ethanol, to eliminate olfactory cues.

#### 2.3.1. Pup retrieval test

On PN7 each dam was removed from their cage for approximately 10 mins, while the pups underwent USV testing. The dam was then returned to her cage, with 5-6 of its pups scattered through three corners of the cage. The dam was video recorded for 10 mins. If the dam failed to return all pups to the nest during this time, the test was stopped and the pups were returned to their nest (Burne et al., 2011; Slamberová et al., 2005). The recorded videos were assessed to determine the time taken for the dam to a) make first contact with her pup (sniffing), b) hold the first pup in their mouth, c) successfully return the first pup to the nest, and d) the time taken to successfully return all 5-6 pups to their nest.

#### 2.3.2 Ultrasonic Vocalisation (USV)

Communication between rats can occur through the emission of Ultrasonic Vocalisations (USVs) (Simola, 2015). Separation of pups from their dam or littermates can result in the rat pups emitting 40 kHz USVs, also regarded as distress calls, due to assessing the situation as threatening and stressful (Simola, 2015). USVs of all pups were recorded on PN7 and PN14 using ultrasound microphones connected to the UltraVox XT system based on previous literature (Potasiewicz et al., 2020). Up to four rat pups at a time were removed from their mother and littermates and placed separately in a clean beaker inside a Styrofoam box, containing a USV microphone to record the rats’ vocalisations. Following three minutes of recording the USVs, the pups were returned to their home cages. Detector outputs were analysed with UltraVox XT (3.0.80) software (Noldus Information Technology, The Netherlands) (Ali et al., 2019). To detect valid calls and enable differentiation of USVs from the background noise, the particular criterion utilised by Ali et al (2019) was applied which involved a minimum amplitude of 40, 20-70 kHz frequency range, 20 millisecond minimum duration with 10 millisecond time gaps. Total number, mean amplitude and duration of 40 kHz vocalisations were measured as per previous studies (van Zyl, Dimatelis and Russell, 2014; Rao and Sadananda, 2015; Ali et al., 2019; Potasiewicz et al., 2020; Lenell et al., 2021). USVs were recorded from a total of 264 pups across 29 litters including 45 WIS-VEH, 91 WKY-Veh and 45 WKY-Sert, and 83 WKY-Ex.

#### 2.3.3. Open Field Test (OFT)

The OFT was used to examine locomotor activity and anxiety-like behaviour. The OFT apparatus consists of a 60cm×60cm square arena (black matte) with 40cm high walls with an even low light of ∼20 lux (Millard et al., 2019; Weston-Green et al., 2011). A single rat was placed in the centre of the arena and behaviour was video recorded for 15 mins. All recordings were analysed using video tracking software (Ethovision, Noldus Information Technology), with the following parameters measured: total travelled distance (in cm) (as a measure of locomotor activity), time spent in the centre of the arena and time spent in the corners (in seconds) (as measures of anxiety-like behaviour).

#### 2.3.4. Novel Object Recognition Test (NOR)

The NOR test is used to examine recognition memory, which is impaired in depression (Kersten et al., 2019; Mathiasen and DiCamillo, 2010). The NOR test arena was the same area as the OFT arena that the rats were previously exposed to. Each rat was allowed to familiarise to two plastic blocks, positioned in the upper half of the arena, for 10 mins. The rat was returned to its home cage and after 60-90 min, the rat was returned to the arena but with one of the plastic blocks replaced with a novel (different-shaped) plastic block. The rat was left to explore the objects for 5 min and was video recorded. The positioning of the new object was altered between the right and left sides, to make sure that the results were not biased based on any side preference of the rats. Interaction time with both the novel and familiar objects were recorded. As rats have a natural tendency to explore new objects, low novel object discrimination is an indicator of impaired recognition memory (Lum et al., 2021; Osborne et al., 2017). Object interaction time was defined as time spent nosing, sniffing, licking or touching the objects with forepaws. A discrimination ratio for each dam was calculated as *T*_N_ / *T*_TOT_; where (*T*_N_ = novel object exploration time, *T*_TOT_ = total object exploration time). A discrimination ratio score of 1 indicated a preference for the novel object, whereas a score closer to 0 indicated a greater preference for the familiar object (Lum et al., 2021; Osborne et al., 2017).

#### 2.3.5. Elevated plus maze (EPM)

The EPM was used to assess anxiety-like behaviour as previously described (Millard et al., 2019). The maze apparatus consisted of 2 closed arms, and 2 open arms, with respective dimensions of 50cm×7cm×30cm (length×width×height) and 50cm×7cm×1cm. The apparatus was arranged around a centre space, with the arms orthogonal to each other. The EPM apparatus was elevated 60cm above the ground and there had a light intensity of ∼100 lux across the open arms. Each dam was placed in the middle of the EPM apparatus and video recorded for 7 mins. Videos were subsequently analysed using Ethovision software to assess time spent in the open arms of the EPM versus the closed arms, corresponding to decrease or increase anxiety-like behaviours as the animal engages in or refrains from exploratory behaviour.

#### 2.3.6. Sucrose Preference Test (SPT)

The SPT was conducted to measure anhedonia (lack of pleasure) as a symptom of depression, similar to previous studies (Brenes Sáenz et al., 2006). Anhedonia would reduce preference for the palatable sweetened water and is an important indicator of depressive-like symptoms (Belovicova et al., 2017). Each dam was single housed to ensure accurate readings of the intake of water vs sucrose, with 2 bottles of tap water for 24 h to habituate the dam to their new cage. After the acclimatisation, one water bottle was replaced with a 3% sucrose solution for 24 h (Hernández et al., 2014). Total water intake was recorded on day 1 of the test and water and sucrose intake were recorded on day 2 by weighing the bottles. Preference for the sucrose solution was measured based on sucrose vs water volume consumed, as a percentage of total fluid consumption over 24 h. One sample t-tests were used to determine whether the rats had a preference for bottles on a particular side (with 50% indicating no preference). While the mean intake from the right side was 60-65%, this was not statistically significant (*p*>0.05). However, to account for any possible left/right-sided preference, the water and sucrose bottles in the SPT were randomly allocated to the left or right side. Following the test period, rats were returned to their original housing conditions.

### 2.4. RNA extraction and quantitative RT-PCR (q RT-PCR)

Dams were euthanized 48 h after their final behavioural test by carbon dioxide (CO_2_) asphyxiation, and their brains were dissected, weighed and stored at -80°C for molecular analyses. The PFC (Bregma level +3.72mm to +2.52mm) and ventral hippocampus (Begma level -2.92mm to - 6.00mm) were dissected according to a standard rat brain atlas (Paxinos and Watson, 2007). RNA extractions and RT-PCR was performed as previously described (Maksour et al., 2024). Briefly, total RNA was extracted using the PureLink RNA Mini Kit (Thermo Fisher Scientific, 12183025) then quantified via a Nano Drop 2000 Spectrophotometer (Thermo Fisher Scientific). The acceptable absorbance ratio of the A260/280 was from 1.9 to 2.1. 0.9 µg of extracted RNA was converted to cDNA using the iScript gDNA Clear cDNA Synthesis Kit (Bio-Rad, 1725035), then qPCR was performed using PowerUp SYBR Green Master Mix (Thermo Fisher Scientific, A25778) with 400 nm of each primer for*, Dnmt1, Dnmt3a, Grin1, Grin2a, Grin2b* (Table S1). We focused on DNA methylation markers initially, based on previous work using fluoxetine (Gemmel et al., 2016) and glutamatergic markers based on increasing implication of the glutamatergic system in depression and its treatments (Khoodoruth et al., 2022). Primer annealing temperatures were optimized by a serial dilution standard curve of each primer, the range of 85-110% efficiency was considered acceptable. RT-qPCR was performed using the QuantStudio 5 Real-Time PCR System (Thermo Fisher Scientific). Each sample was run in triplicate; each reaction contained 20ng of cDNA template in a final reaction volume of 20 µL. The cycling parameters were: 50°C for 2 min, 95°C for 2 min, then 40 cycles of 95°C for 1 s and 30 s for the annealing/extending at 60°C. The reaction was followed by conditions for melt curve analysis: 95°C for 15s, 60°C for 1 min and 95°C for 15s. ΔCt values of the *Dnmt1*, *Dnmt3a, Grin1, Grin2a, Grin2b,* mRNA quantities were obtained by normalisation to the average of reference genes including, *B2m*, *Ubc*, *Gusb*, *Ppia,* and *Gapdh.* Data are presented using the 2-ΔCt calculation to yield relative gene expression values.

### 2.5. Statistical analyses

All statistics were performed by IBM SPSS, version 29. One dam from the WKY-Sert group was excluded from the data analysis due to the cannibalism of her pups. Boxplots were used to identify statistical outliers (data points located further than three times the interquartile range). The average daily running distance in the WKY-Ex group was compared between days 1-6, 7-12, and 13-18 using a repeated-measures ANOVA. One-Way ANOVAs were used to compare the weight differences before mating, then repeated-measures ANCOVA, covarying for litter size, to analyse time*group interaction on dam weight gain/loss during and after pregnancy. If/when there was a significant time*group interaction, one-way ANCOVAs covarying for litter size were used to assess the impact of the intervention (sertraline, exercise, or vehicle) on dam body weight gain/loss at each specific stage (GD0-7, GD8-14, GD15-PN0, PN1-7, and PN8-14) separately. ANOVAs/ANCOVAs were used to compare litter size, offspring average body weight at PN7 and PN14 (covarying for litter size), pups’ brain weight and dams’ brain weight (covarying for body weight), parameters of the behavioural tests (OFT, EPM, NOR and SPT), the time the dam took to make first contact (sniffing) with a pup in the pup retrieval test, and the mRNA levels. Post hoc tests were used to detect significance differences between the groups. Chi-square tests were used to examine the effects of treatment on measures in the pup retrieval test including whether there were differences in the number of dams that interacted with the pups via a) grabbing at least one pup in its mouth, b) returning at least 1 pup to the nest, and (c) whether there were differences in the number of dams that completed the test (i.e. returned all pups to the nest). USVs at PN7 were examined using ANOVA. USVs at PN14 were consistently not normally distributed, so were analyzed using Kruskal-Wallis test overall and in male and female pups separately. Mann-Whitney U test was used to examine any differences between male and female pups. Significance was set at an alpha level of *p*<0.05.

## 3. Results

### 3.1. Running wheel data

The total running distance of each dam in the WKY-Ex group ranged from 2000m to 8000m over 18 days, with a median value of 3817m (Figure 2a). The average running distance during gestational days (GD) 1-6 and 7-12 of pregnancy were similar, followed by a decline in the average running distance at GD 13-18 compared to GD 1-6 (-26%; *p*=0.056) and GD7-12 (- 34%; *p*=0.009) (Figure 2b).

**Figure 2:**
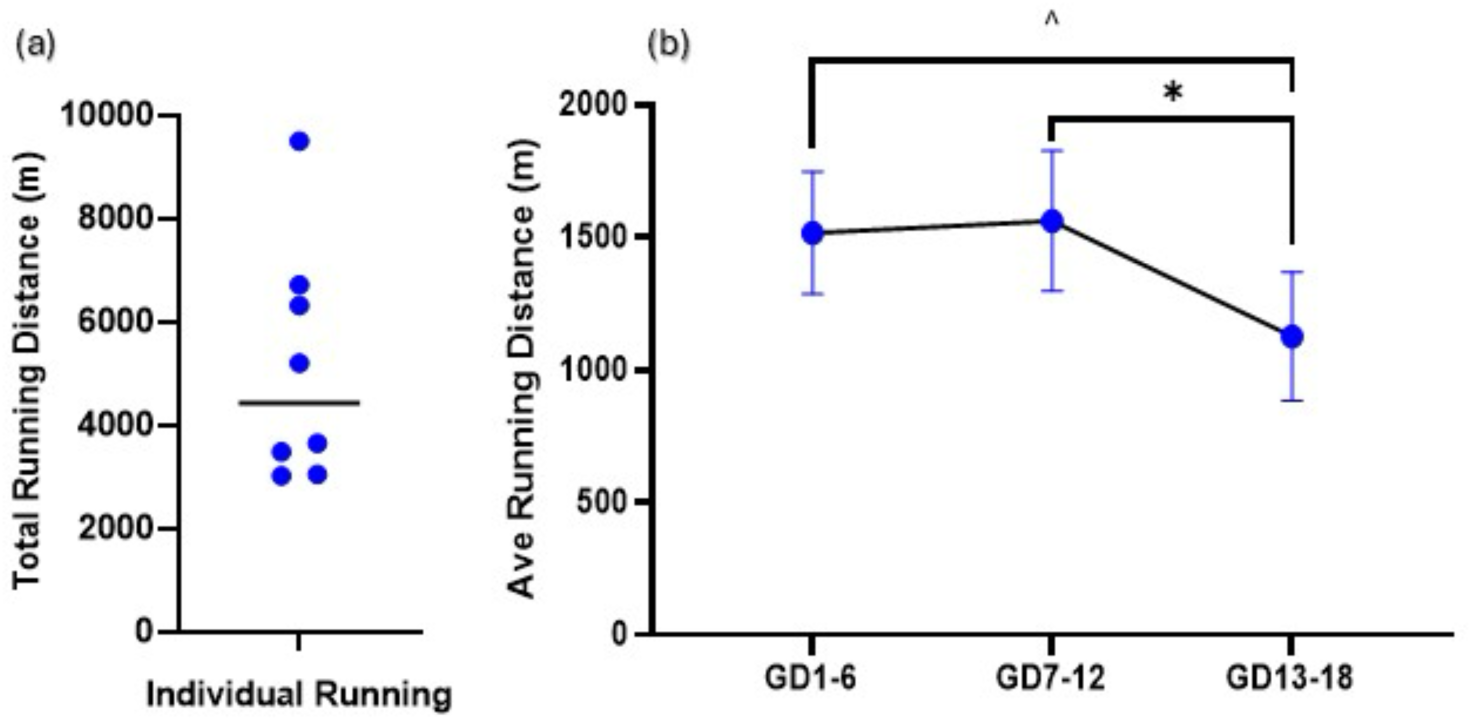
Running wheel outcome for the WKY-Ex group. (a) total running distance undertaken by each dam; the horizontal line represents the median distance; (b) average distance undertaken by the WKY-Ex group through 3 gestational periods (of a period of 6 days each). GD: gestational day; data expressed as mean distance in meters (+ SEM); ^:p=0.056 compared to GD 1-6;*:p<0.05 compared to GD 7-12.

### 3.2. Dam characteristics

#### 3.2.1 Dam body weight

One way ANOVA showed that dam body weight prior to mating was significantly different between the 4 groups (F₁,₂₉=168.745, *p*<0.001). The three WKY groups weighed 31-34% less than the Wis group (*p*<0.001) but there was no difference between the WKY groups (Figure 3.7b). The repeated measures ANCOVA (covarying for litter size) of the percentage of weight gained/lost during pregnancy up to PN14, revealed a significant time effect (F₁,₄=205.597, p<0.001) and a significant interaction of time*group (F₁,₁₂=2.733, p=0.004) between the four dam groups (Figure 3a). Post hoc testing revealed that the percentage of body weight gained and lost in the WKY-Sert group was significantly different compared to the Wis (*p*=0.018) and WKY-Ex (*p*=0.005) groups and a trend towards significance compared with the WKY-Veh group (p=0.08). Specifically, the WKY-Sert group gained 39% less weight between mating and GD7 compared to the Wis (*p*=0.023), WKY-Ex (*p*=0.012) and WKY-Veh (*p*=0.012) groups (Figure 3c). There was no difference in percentage of body weight gain between GD7 and GD14 between the four groups (F₁,₂₈=2.151, *p*=0.121; covarying for litter size) (Figure 3d). However, the percentage of body weight gained from GD14 to the delivery day was significantly different between the 4 groups (F₁,₂₉=3.786, *p*=0.024; covarying for litter size), where all the WKY groups gained 39% less weight compared to the Wis group (*p*<0.05) (Figure 3e).

**Figure 3:**
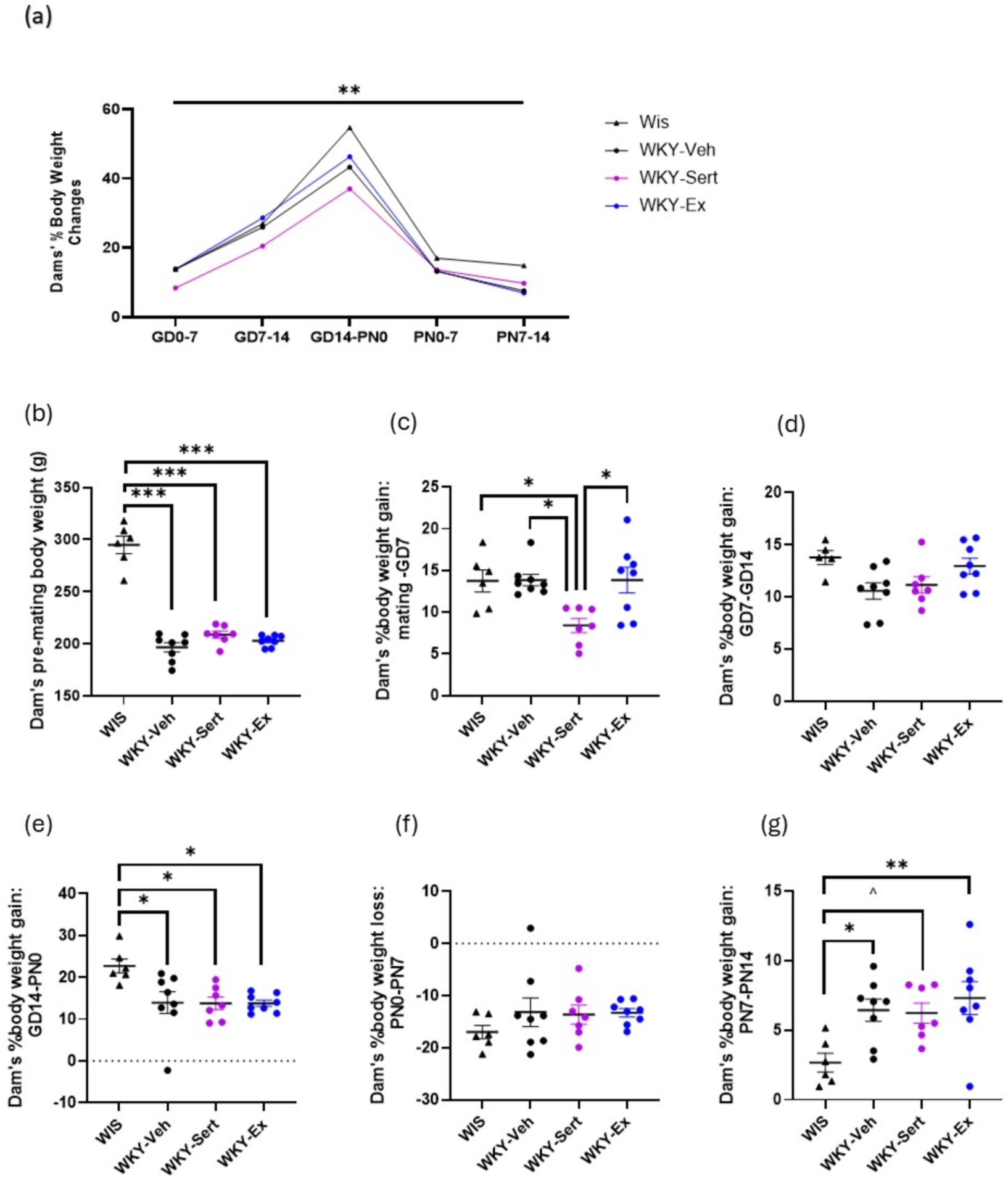
Dam’s Body Weight Changes Across Gestational & Postnatal Period. (a) Summary of dams’ percentage body weight change over the time. (b) Dam’s pre-mating body weight; (c) Dam’s percentage body weight gain from mating to gestational day 7 (GD7); (d) Dam’s percentage body weight gain from GD7 to GD14; (e) Dam’s percentage body weight gain from GD14 to delivery day (PN0); (f) Dam’s percentage body weight loss from PN0 to postnatal day 7 (PN7); (g) Dam’s percentage body weight gain from PN7 to PN14; in Wistar (WIS; n=5-6), Wistar Kyoto Vehicle (WKY-Veh; n=8), Wistar Kyoto sertraline (WKY-Sert; n=7), and Wistar Kyoto exercise (WKY-Ex; n=8) groups. Data are expressed as mean % change weight +/- SEM. GD: gestational day, PN: postnatal day. ^: *p*=0.065, *: *p*<0.05, **: *p*<0.01, ***: *p*<0.001.

The percentage of body weight lost from PN0 to PN7 was not significantly different between the four groups (F₁,₂₉=1.045, *p*=0.391; covarying for litter size) (Figure 3f). There was however a significant difference between the groups with regards to percentage of body weight change from PN7 to PN14 (F₁,₂₉=3.98, *p*=0.02; covarying for litter size), with all WKY groups losing a larger percentage of body weight compared to the Wis group, reaching significance in the WKY-Ex (*p*=0.009) and WKY-Veh (*p*=0.039) and approaching significance in the WKY-Sert group (*p*=0.065) (Figure 3g).

### Litter characteristics

#### 3.2.1. Litter size

Litter size was significantly different between the four groups (F_1,29_=6.062, *p*=0.003); the WKY-Sert group showed a 43% and 38% smaller litter size compared to Wis (*p*=0.003) and WKY-Ex (*p*=0.012) groups, respectively (Figure 4a). The ratio of females to males in each litter was not significantly different between the groups (F₁,₂₉=1.745, p=0.183) (Figure 4b).

**Figure 4:**
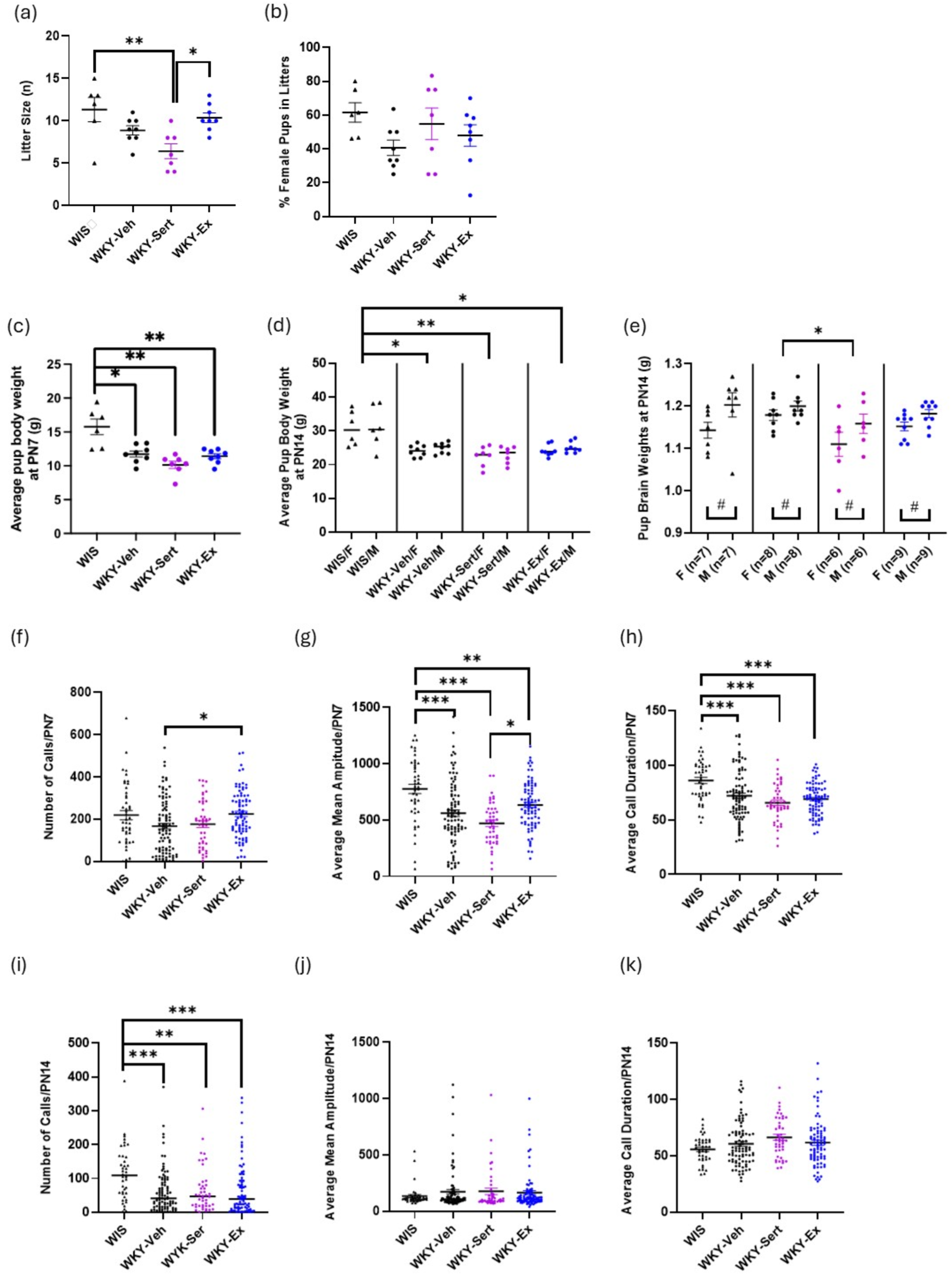
Litter characteristics. (a) Litter size (number of pups/litter), (b) Percentage of female pups in each litter, (c) Average pup body weight on postnatal day 7 (PN7), (d) Average pup body weight on postnatal day 14 (PN14) of both sexes, in Wistar (WIS; n= 6), Wistar Kyoto Vehicle (WKY-Veh; n=8), Wistar Kyoto sertraline (WKY-Sert; n= 7), and Wistar Kyoto exercise (WKY-Ex; n=8 groups). (e) Pup brain weights at PN14 of both sexes, in WIS; n= 14, WKY-Veh; n=16, WKY-Sert; n= 12, and WKY-Ex; n=18 groups. USVs at PN7: (f) Number of calls, (g) Average Mean Amplitude, (h) Average call duration. USVs at PN14: (i) Number of calls, (j) Average Mean Amplitude, (k) Average call duration, in WIS; n=41-45, WKY-Veh; n=89-94, WKY-Sert; n=44-45, and WKY-Ex; n=81-83 groups. USV: Ultrasonic vocalizations, WIS: Wistar, WKY-Veh: Wistar Kyoto-Vehicle, WKY-Sert: Wistar Kyoto-Sertraline, WKY-Ex: Wistar Kyoto-Exercise. Data expressed as mean + SEM, *: p<0.05, **: p<0.01, ***: p<0.001.

#### 3.2.2 Pup characteristics

Pup average body weight at PN7 and PN14 differed between the four groups after covarying for litter size (PN7: F_1,29_ =8.604, *p*<0.001; PN14: F_1,29_=5.472, *p*=0.005). At PN7, the average pup body weight in all WKY groups was significantly less than the Wis group; WKY-Veh and WKY-Ex groups weighed 25-27% less and WKY-Sert weighed 36% less than the Wis group at PN7 (*p*≤0.001; Figure 4c). Similarly, at PN14, the average body weight of all WKY groups was less than the Wis group; the WKY-Veh and WKY-Ex groups weighed 20-21% less (p<0.01), and WKY-Sert weighed 27% less than the Wis group (*p*<0.001) (Figure 4d). There were no significant differences in body weight between the WKY groups. When examining PN14 body weights in male and female offspring separately, WKY offspring of both sexes consistently weighed less than the Wis-group (male: F_1,29_=4.631, *p*<0.011; female: F_1,29_=6.042, *p*<0.003).

#### 3.2.3 Pup Ultrasonic Vocalisations

At PN7, one-way ANOVA showed a significant difference between the four groups in the number of calls (F_3, 264_ = 4.021, *p*=0.008). Post hoc analysis showed that the WKY-Ex group had a 35% higher number of calls compared to WKY-Veh (*p*=0.013) (Figure 4f). The average mean amplitude was significantly different between the four groups (F_3, 264_ = 12.632, *p<*0.001). Post hoc analysis showed that the WKY groups displayed an 18-39% reduction compared to the Wis group (*p*<0.005) and WKY-Ex pups displayed a 34% increase in mean amplitude compared to WKY-SERT (p=0.003) (Figure 4g). Similarly, the average of the call duration was significantly different between the four groups (F_3, 264_=10.594, *p<*0.001). Post hoc analysis showed that the WKY groups displayed a 16-23% reduction compared to the Wis group (*p*<0.001) (Fig. 4h).

At PN14, Kruskal-Wallis showed that the number of calls was significantly different between the groups (χ^2^(3)=19.167; p<0.001), with a reduction in WKY-VEH (-46%; p=0.001), WKY-SERT (-43%; p=0.009) and WKY-EX (-41%; p<0.001) groups compared to WIS control (p<0.01) (Figure 4i). There was however no significant difference between groups in average mean amplitude (χ^2^ (3) =1.035; p=0.793) (Figure 4h) or average call duration (χ^2^(3)=6.952; p=0.073) (Figure 4j). When male and female groups were analysed separately, similar patterns of significance were found (Supplementary materials). There was no difference between male and female offspring, regarding number of calls (*U*=8008.5; p=0.900). However average call duration was 12% longer in males (*U* =9726.5; p=0.007) and average mean amplitude was 43% larger in males compared to female pups (*U* =9411; p=0.029).

#### 3.2.4. Pup brain weight at PN14

When analysing brain weight of male and female pups at PN14, covarying for body weight, there was a significant main effect of group (F₁,₆₀=9.077, *p*<0.001); the brain of pups in the WKY-Sert group weighed 5% less than the WKY-Veh group (*p*=0.031). There was also a significant main effect of sex (F₁,₆₀=6.592, *p*=0.013), where female brains weighed 3.5% less than males, but there was no significant interaction between group and sex (F₁,₆₀=0.231, *p*=0.875) (Figure 4e).

### 3.3. Dam behaviour

#### 3.3.1. Pup retrieval test (PRT)

All dams interacted with their pups by sniffing the pups during the test period, with no differences between the four groups with the time until the first sniffing interaction (F_1,27_=0.138, *p*=0.937). Not all dams completed the other parameters of the test including grabbing at least one pup in their mouth, returning at least one pup to the nest, or returning all the pups into their nest. Accordingly, these parameters were analysed using the Chi-square test (with dams being recorded as ‘completers’ or ‘non-completers’). There was no significant difference between the four groups regarding the number of dams that grabbed at least one pup in their mouth (*p*=0.441) or returned at least one pup to the nest (*p*=0.263). There was, however, a main effect of group with regard to the number of dams that completed the test (i.e., successfully returned all the pups to their nest within the allocated time) (p=0.05). None of the WIS dams finished this task, with code cross-tabulation showing only the WKY-Ex group had a greater percentage of dams that completed the test compared to the WIS group. While it appeared that the WKY-Ex group also performed better than the WKY-Veh and WKY-Sert groups, this did not reach significance, likely due to limited power (Table S3).

#### 3.3.2. Open field test (OFT)

When tested 5 weeks postnatally, distance travelled was significantly different between the four groups (F₁,₂₉=17.111, *p*<0.001); all three WKY groups travelled less than the Wis group (27-41%; *p*<0.001) (Figure 5a). The accumulated time spent in the centre showed a trend towards a significant difference between the four groups (F_₁,₂₇_=2.796, *p*=0.063), with the WKY-Ex group spending 132% more time in the centre than the WKY-Veh (*p*=0.057) (Figure 5b). There was no significant difference between the groups in time spent in the corners of the open field apparatus (F₁,₂₉=.0584, *p*=0.631) (Figure 5c).

**Figure 5:**
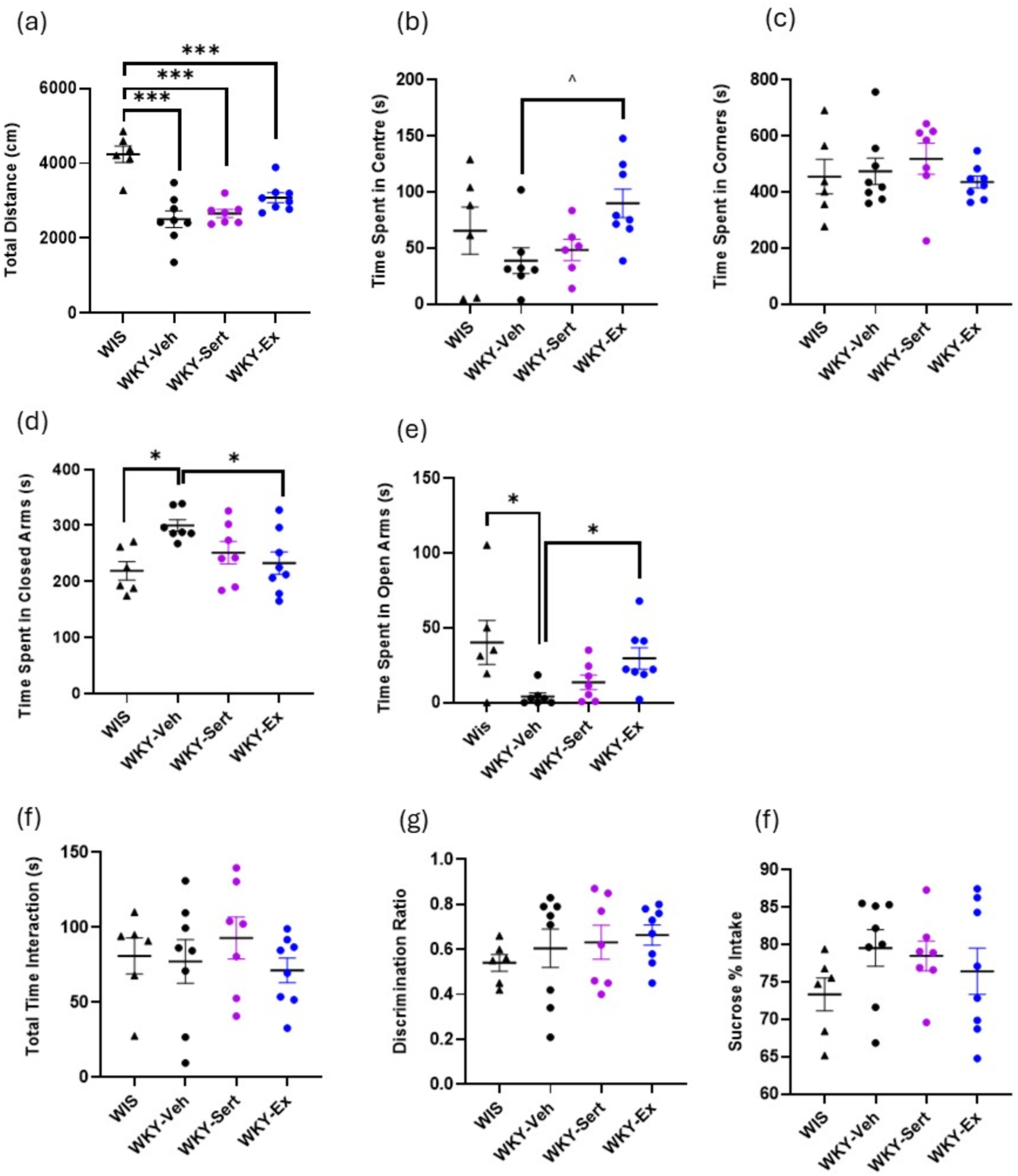
Dams’ Postnatal Behavioural Tests: OFT. (a) total distance (b) time spent in centre, (c) time spent in corners, **EPM** (d) time spent in closed arms, (e) time spent in open arms, **NOR** (f) total time interaction, (g) discrimination ratio, **SPT** (h) sucrose percentage intake over 24 hours, in Wistar (Wis; n=6), Wistar Kyoto Vehicle (WKY-Veh; n=7-8), Wistar Kyoto sertraline (WKY-Sert; n=6-7), and Wistar Kyoto exercise (WKY-Ex; n=8). EPM: elevated plus maze, NOR: novel object recognition OFT: open field test, SPT: sucrose preference test. Data expressed as mean + SEM, *:p<0.05, **p<0.01, ***:p<0.001, ^:p=0.057.

#### 3.3.3. Elevated plus maze (EPM)

Time spent in the closed arms (F₁,₂₈=3.933, *p*=0.020) and open arms (F₁,₂₉=4.014 *p*=0.021) of the EPM were significantly different between the four groups. Post hoc analysis showed that the WKY-Ex group spent 22% less time in the closed arms compared to WKY-Veh (*p*=0.047) and that the WKY-Veh group spent 37% more time in the closed arms compared to the Wis group (*p*=0.022) (Figure 5d). The WKY-Veh group spent 90% less time in the open arms compared to the Wis group (*p*=0.038), and the WKY-Ex spent 620% more time in the open arms compared to the WKY-Veh group (*p*=0.027) (Figure 5e).

#### 3.3.4. Novel object recognition (NOR)

There were no group differences between total object interaction time (F₁,₂₉=0.546, *p*=0.655) or discrimination index (F₁,₂₉=0.557, *p*=0.649), (Figure 5f).

#### 3.3.5 Sucrose preference test (SPT)

There was no significant difference in the percentage of sucrose intake between the four groups (F₁,₂₉=1.058, *p*=0.385, covarying for dam body weight) (Figure 5g).

### 3.4. Dam brain weight

Dam brain weights were significantly different between the four groups after co-varying for dam body weight (F₁,₂₇=5.999, *p*=0.004); the WKY-Veh, WKY-Sert and WKY-Ex brains weighed 9, 12 and 11 % more than the Wis brains, respectively (*p*<0.001), but there were no differences between the three WKY groups (Figure 6a).

**Figure 6:**
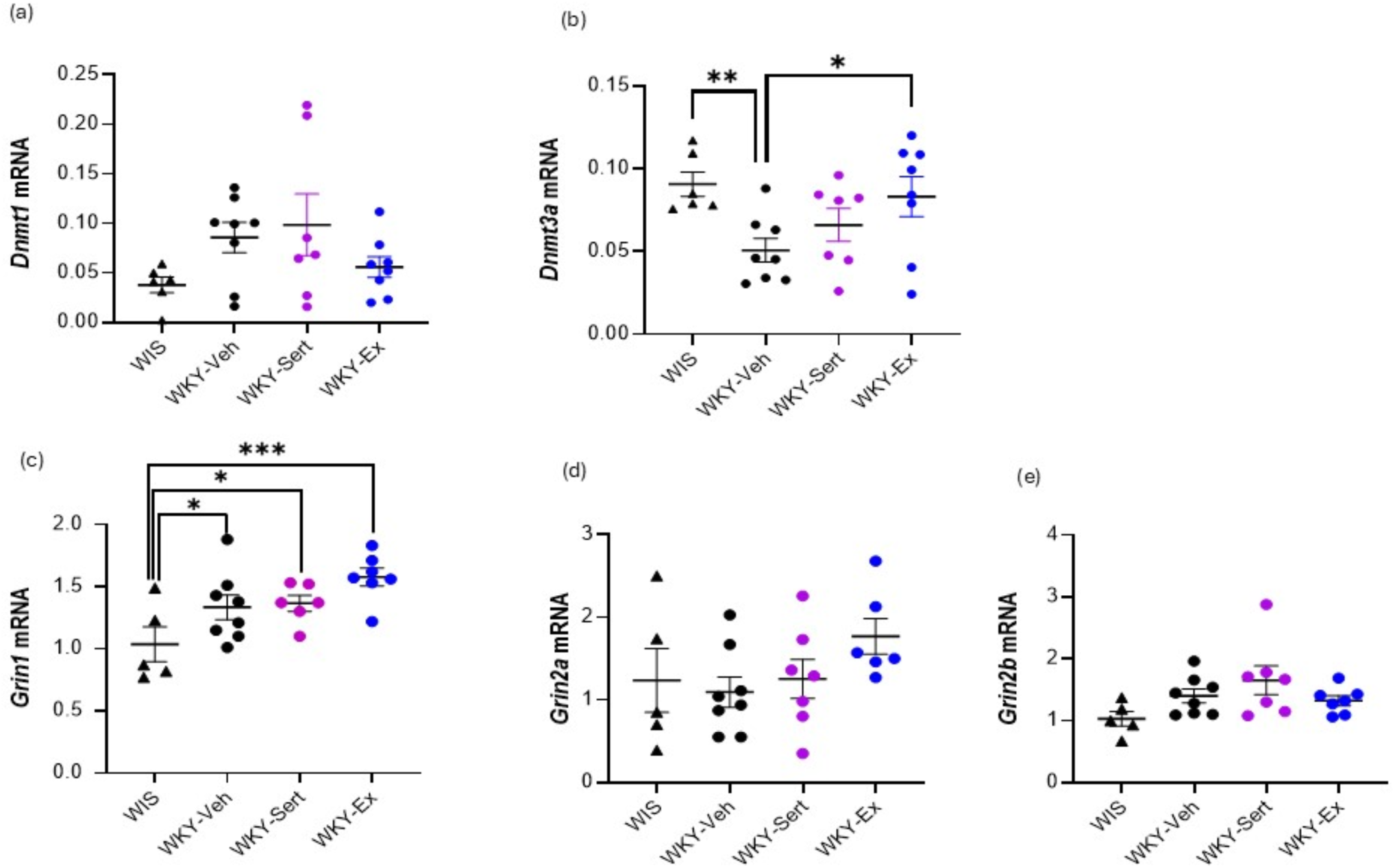
Dam mRNA Gene Expression Levels. (a) *Dnmt1,* (b) *Dnmt3a,* (c) *Grin1,* (d) *Grin2a,* (e) *Grin2b* relative expression (2 ^ΔCt^) in the PFC in Wistar (Wis; n=5-6), Wistar Kyoto Vehicle (WKY-Veh; n=8), Wistar Kyoto sertraline (WKY-Sert; n=6-7), and Wistar Kyoto exercise (WKY-Ex; n=6-8) groups. Data expressed as mean ± SEM, *:*p*<0.05, **:*p*<0.01, ***:*p*<0.001.

### 3.5. Dam Gene Expression Levels

The *Dnmt3a* mRNA level was significantly different between the four groups (F_3, 29_=3.439, *p*=0.032). Post-hoc analysis showed that the WKY-Veh group had 44% less *Dnmt3a* mRNA expression compared to the Wis group (*p*=0.009), and the WKY-Ex reported a 64% increase in *Dnmt3a* mRNA expression compared to the WKY-Veh group (*p*=0.019) (Figure 6b). The *Dnmt1* mRNA level however did not show any significant difference in the PFC between the four groups (F_3, 29_=2.063, *p*=0.131) (Figure 6a). A one-way ANOVA revealed that *Grin1* expression was significantly altered by treatment (F_3, 26_=4.93; p=0.009). Post-hoc analysis revealed a significant 29%, 32%, 52% increase in *Grin1* in WKY-Veh (p=0.041), WKY-SERT (p=0.034) and WKY-Ex (p<0.001) compared to WIS (Figure 6c). There were no significant differences in *Grin2a* (F_3, 26_=459; p=0.713) or *Grin2b* (F_3, 26_=2.564; p=0.079) between the groups (Figure 6d-e). Gene expression in the ventral hippocampus was not significantly different between groups (Table S2).

## 4.0 Discussion

In this study, we showed that maternal sertraline treatment, at 20mg/kg/day, during pregnancy and the early postnatal period, in the WKY rat, resulted in less weight gain during pregnancy and a smaller litter size, with pups having slightly smaller brain weights, compared to vehicle controls. Sertraline treatment however did not alter dam postnatal measures of activity, anxiety, anhedonia or recognition memory in the dams, or *DNMT* or *Grin* gene expression in the PFC. In contrast we showed that voluntary exercise during pregnancy, did not impact dam body weight during pregnancy; it did result in more 40kHz calls in pups at PN7, but did not alter other litter characteristics. Importantly, voluntary exercise during pregnancy reduced measures of postnatal anxiety in the dam, accompanied by an increase in cortical *DNMT3a* gene expression.

### 4.1 Maternal sertraline treatment in the WKY rat does not impact postnatal activity, anxiety, anhedonia or cognitive behaviours

The present study did not show any impact of maternal sertraline treatment on postnatal measures of activity, anhedonia, anxiety or recognition memory, when tested in postnatal week 5, or on maternal behaviour as measured by the PRT. Currently, there is no published literature on the effect of maternal sertraline treatment on postnatal behaviour. However, Amani et al., (2021) reported that 10 and 25mg/kg (but not 5mg/kg/day) of maternal fluoxetine treatment (GD10-PN20) reversed the effect of prenatal stress on anxiety- and depressive-like behaviours through the EPM and forced swim test, when tested in the first postnatal week (Amani et al., 2021). Similarly, Salari et al (2016) reported that maternal fluoxetine (8mg/kg/day, GD10 to PN20) reversed the effect of gestational-stress on depressive-like behaviour, as measured by the SPT and forced swim test, in the first postnatal week (Salari et al., 2016). In contrast, postpartum fluoxetine (10 mg/kg/day, PN2-25) exacerbated depressive- and anxiety-like behaviours as measured via the forced swim test and novelty suppressed feeding test, in postnatal week 4, in both control and postpartum induced stress dams (Gobinath et al., 2018). Similarly, Pawluski et al., (2012) found that postpartum fluoxetine treatment (5 mg/kg/day, on PN0-28) increased anxiety-related behaviour (in the elevated zero maze) but this was only in the control (non-stressed) rats and not in the gestationally stressed Sprague-Dawley dams (Pawluski et al., 2012). Collectively this might suggest that maternal fluoxetine treatment has a greater ability to modulate maternal behaviour than sertraline (at the dose used in this study), and that the period of exposure may influence the outcome. In the present study, we measured the delayed effect of sertraline at postnatal week five, which was three weeks after maternal sertraline treatment had stopped. Had we measured behaviours earlier in the postnatal period, or extended sertraline treatment, therapeutic effects may have been seen. Furthermore, implementing a wider test battery would better characterise the impact of sertraline on these behavioural domains. In the clinical scenario, it is unlikely that sertraline or SSRI treatment more generally, would be stopped abruptly at this early postnatal stage, which may have contributed to the findings in this study. In addition, the WKY strain, used in the present study, is known to model elements of treatment resistant depression (Millard et al., 2020) which may make it particularly difficult to see improvements in relevant behaviours following SSRI treatment. The available evidence shows that WKY rats display a hypo-serotonergic phenotype (low serotonin transporters) mainly in cortex and the hippocampus brain regions which might explain the strains antidepressant resistance (Coplan et al., 2014; Paré and Tejani-Butt, 1996).

In line with our findings showing no effect of sertraline on maternal behaviour, we also observed that maternal sertraline treatment did not have any significant impact on *Dnmt or Grin* mRNA levels. While no study has assessed the impact of maternal sertraline on the maternal brain, our finding was consistent with Gemmel et al., (2016) who reported that postpartum fluoxetine (5 mg/kg/day, PN1-21) in a gestational stress Sprague Dawley model did not influence DNMT3a protein level in the PFC, in both the stress and control groups (Gemmel et al., 2016). Sertraline treatment during pregnancy (2.5 or 10mg/kg/day) has previously resulted in fewer Ki67 and doublecortin immunoreactive neurons in the hippocampus (Pawluski et al., 2020). Similarly, Gemmel et al (2016) showed a decrease in DNMT3a density in the hippocampus in fluoxetine exposed dams, suggesting the hippocampus may be more sensitive to SSRI induced changes however we found no evidence of gene expression changes in the hippocampus following sertraline treatment.

Despite no sertraline-induced changes in dam outcomes in this study, our findings suggest that maternal sertraline treatment may have mild effects on early offspring development, as we show that the sertraline group gained the least weight during pregnancy, delivered a smaller litter, the pups of which had slightly smaller brain weights, although there was no change in pup body weight. Similar findings have been reported showing a reduction in body weight in rat dams at GD20 following sertraline (20mg/kg/day) from GD13-GD20, compared to non-sertraline treated controls (Lozano et al, 2021) and no changes in offspring body weight at PN7 or PN14 (Moura et al., 2023), although Moura et al (2023) did report a small reduction in body weight a PN1. In contrast, Kummet et al (2012) reported a reduction in the body weight of pups when a low dose of sertraline (5 mg/kg) was administered *directly* to the pups from PN1-14. This highlights the difference between gestational and direct postnatal exposure and may reflect the minimal levels of sertraline transmitted to infants via breast-feeding during the prenatal period (Kristensen et al., 1998; Retz et al., 2008; Anderson and Sauberan, 2016; Pogliani et al., 2019). In contrast, previously our lab showed that maternal fluoxetine treatment (10 mg/kg, GD0-PN14) did not impact dam body weight or litter size in Sprague Dawely and WKY rats (Millard et al., 2019) but did result in lower body weight in male and female offspring at PN7 and PN14 (Millard et al., 2019). These findings point towards different outcomes between the two SSRIs, which could be associated with the longer half-life of fluoxetine or the greater transfer from the dam to the infant via breast-feeding (Pogliani et al., 2019). It will be important to conduct further analysis of the sertraline exposed offspring from this study to determine if there are any longer term behavioural or molecular impacts of these perinatal changes.

### 4.2. Voluntary exercise during pregnancy reduced postnatal anxiety-like behaviour in the dam and increased cortical Grin1 gene expression

The present study revealed that maternal voluntary exercise reduced postnatal anxiety-like behaviour through the OFT and EPM. This is the first study, to our knowledge, that has examined the effect of exercise during pregnancy, on postnatal anxiety (as well as depressive and cognitive) behaviours. Previous work has however measured maternal care related behaviours; Gobinath et al., (2018) previously reported in a postpartum depression model that maternal exercise (8 week free running wheel: pre-mating, through gestation, and 2 weeks after delivery) could not reverse the postnatal corticosterone treatment-induced reduction in maternal care (time spent in nursing and time spent off the nest) (Gobinath et al., 2018). Interestingly, in this same model, Gobinath et al., reported that in high running dams, but not the low running dams, fluoxetine treatment significantly increased time spent nursing and reduced time spent off nest, suggesting the higher voluntary exercise positively influenced efficacy of fluoxetine in maternal care (Gobinath et al., 2018c). While our study was not powered to distinguish between high and low runners, behavioural data in the WKY-EX group, for high and low runners is presented in Supplementary Table 3, showing high runners spent significantly more time with the novel object than low runners. The work of Gobinath also does raise the question of whether sertraline combined with exercise would have an interactive effect in pregnancy and something that should be considered in future studies. While there are limited studies examining the effects of maternal exercise on postnatal behaviour, there is a wealth of data in non-pregnant rodent models showing the positive impact of exercise on anxiety-like and depressive-like behaviours (Binder et al., 2004; Fulk et al., 2004; Fuss et al., 2010; Patki et al., 2014; Uysal et al., 2014; Yüksel et al., 2019). Our findings suggest the benefits exercise extend to pregnancy. Our review of clinical studies has provided preliminary evidence that maternal exercise could potentially improve anxiety and depression symptoms during and after pregnancy (Jarbou and Newell, 2022). Taken together, this suggests future studies investigating the effects of exercise in pregnancy in the human population are warranted.

In addition to the observed behavioural benefits of exercise, we also show reduced cortical *Dnmt3a* mRNA (but no change in *Dnmt1)* in the WKY-Vehicle group compared to Wistar controls and a subsequent increase in *Dnmt3a* mRNA in the WKY exercise group suggesting DNA methylation may contribute to the beneficial effects of maternal exercise. Evidence indicates that DNA methylation plays a key role in how exercise influences gene expression (for review see: Fernandez et al, 2018) and our findings indicate, for the first time, this extends to exercise in pregnancy. Further studies, using methyl-ATAC-seq or similar to determine whether DNMT changes reflect DNMA methylation to specific genes is warranted. We did also show that all WKY groups presented with increased *Grin1* gene expression; it appeared that sertraline or exercise during pregnancy was not above to lower these levels.

In addition to the positive outcomes that exercise produced in the dams, it did not influence dam or litter characteristics, indicating the safety/tolerability of exercise interventions in pregnancy. WKY-Ex offspring, did however, produce more 40kHz USVs at PN7, but not at PN14, suggesting this effect was transient. While an increase in 40kHz calls has been suggested to be associated with greater distress or anxiety (Kim et al., 2010; Simola, 2015), these isolation induced calls have also been suggested to represent enhanced communication with the dam for retrieval (Ehret, 2005; Shekel et al., 2021). This corresponds to better performance by the WKY-Ex dams in the PRT, suggesting it could be an effect driven by maternal behaviour. Considering the WKY groups generally showed reductions in several USV parameters compares to WIS controls, this increased call number at PN7 could represent an exercise induced adaptation to improve social communication.

A consideration with these findings in the WKY-Ex group is that they may not be driven by maternal exercise, but rather by the environmental enrichment that was associated with the daily change to the exercise cage environment. Studies have shown that environmental enrichment has the ability to improve anxiety-like and depressive-like behaviour and spatial functions in rats (Burrows et al., 2011; Harati et al., 2013; Jin et al., 2017; Leggio et al., 2005). In the present study, rats were housed in individually ventilated cages (IVCs) throughout the study and while the majority of the time they were housed in pairs to ensure opportunities for adequate social interaction, inter-cage olfactory communication is very limited in IVCs (Guerra et al., 2021; Hawkins et al., 2023). However the exercise group were transferred daily to open top cages during their running protocol, which may have allowed for greater social (olfactory) cues. Furthermore, the additional handling required during this period may have contributed to reduced stress and anxiety in this group (Costa et al., 2012).

### 4.3. Conclusion

In conclusion, we have shown for the first time that voluntary (running) exercise during pregnancy in the WKY rodent model reduces anxiety-like behaviour in the dam at five weeks postnatal. This was accompanied by increased *DNMT3a* gene expression in the PFC suggesting this region may be sensitive to methylation changes following maternal exercise. In contrast, treatment with sertraline (at 20mg/kg/day) did not impact these behaviours. It may be that the effects of sertraline would have been seen had treatment to the dams been maintained. Sertraline did however appear to have mild fetal effects, as seen through stunted dam body weight growth early in pregnancy, smaller litter sizes and slightly smaller offspring brain weights, which should be investigated further. Overall, our findings suggest a long-term beneficial effect of exercise during pregnancy on affective behaviour and supports future studies examining the effects of exercise in antenatal depression and anxiety in the human population.

## Declaration of generative AI in scientific writing

Generative artificial intelligence (AI) and AI-assisted technologies were not used in the writing process.

## CRediT authorship contribution statement

*Noor S. Jarbou:* Investigation; Methodology; Formal Analysis; Visualization; Writing – Original draft; Writing – review and editing; *Olivia Mairinger:* Methodology, investigation, writing – original draft; *Elise Kulen:* Investigation; Writing – review and editing; *Lucas Mushawar:* Investigation; Writing – review and editing; *Samara Walpole:* Investigation; Methodology; Formal data analysis; Writing – review and editing; *Jasmine Matthews:* Investigation; Formal data analysis; Writing – review and editing; *Simon Maksour:* Investigation; Methodology; Writing – review and editing; *Katrina Green:* Resources; Writing – review and editing; *Mirella Dottori:* Supervision; Resources; Writing – review and editing; *Kelly A. Newell:* Conceptualization; Methodology; Formal analysis; Investigation, Resources, Writing – review and editing, Supervision, Funding acquisition.

