## Supplementary Data for "THE EFFECT OF SERTRALINE AND VOLUNTARY EXERCISE DURING PREGNANCY ON LITTER CHARACTERISTICS AND POSTNATAL AFFECTIVE BEHAVIOUR IN RAT DAMS"

### Supplementary materials

**Table S1:** Primers used for RT-PCR.

| **Target** | **Forward primer (5' – 3')** | **Reverse primer (5' – 3')** |
|---|---|---|
| *Dnmt1* | AGAGACCAGGATAAGAAACG | TTACTCGTTCAGGTTTCTCC |
| *Dnmt3a* | AATAGCCAAGTTCAGCAAAG | AAACACCCTTTCCATTTCAG |
| *Grin1* | AAGGAGAATATCACTGACCC | TACTTAGAAGACATCAGCACC |
| *Grin2a* | GATCAACAATTCAACCAACG | AGACCACTTCACCTATCATTC |
| *Grin2b* | GTTTAACAACTCCGTACCTG | TCTGGAACTTCTTGTCACTC |
| *Gusb* | GATTCAGATATCCGAGGGAG | CGATGACCACAATTCCATATC |
| *Ppia* | GTGTTCTTCGACTCACG | AAGTTTTCTGCTGTCTTTGG |
| *Gapdh* | GCCTTCCGTGTTCCTACC | CCTGCTTCACCACCTTCTT |

#### USV results in female pups at PN14

At PN14, female rat pups showed a significant difference between groups in number of calls ($\chi^2$ (3)=9.171; p=0.027), with post-hoc showing a significant reduction in the three WKY groups compared tothe WIS group (p<0.05). There was however no significant difference between groups in average call duration ($\chi^2$ (3)=4.483; p=0.214) or average mean amplitude ($\chi^2$ (3)=1.566; p=0.667)

#### USV results in male pups at PN14

At PN14, male rat pups showed a significant difference in the number of calls between groups ($\chi^2$ (3)=13.142; p=0.004), with post-hoc showing a significant reduction in all WKY groups compared to the WIS group (p<0.01). There was however no significant difference between groups in average call duration ($\chi^2$ (3)=5.053; p=0.168) or average mean amplitude ($\chi^2$ (3)=3.789; p=0.285).

**Supplementary Table 2: Gene expression in the ventral hippocampus**

| | ***WIS*** | ***WKY-Veh*** | ***WKY-Sert*** | ***WKY-Ex*** | ***p-value*** |
|---|---|---|---|---|---|
| *Grin1* | 1.033± | 1.180± | 1.025± | 0.791± | 0.400 |
| | 0.139 | 0.295 | 0.158 | 0.112 | |
| *Grin2a* | 1.085± | 1.324± | 1.474± | 0.961± | 0.129 |
| | 0.184 | 0.327 | 0.090 | 0.123 | |
| *Grin2b* | 1.117± | 1.452± | 1.537± | 1.104± | 0.367 |
| | 0.163 | 0.458 | 0.114 | 0.170 | |
| *Dnmt1* | 1.038± | 1.012± | 0.977± | 1.145± | 0.775 |
| | 0.125 | 0.082 | 0.109 | 0.139 | |
| *Dnmt3a* | 1.016± | 0.971± | 0.942± | 1.241± | 0.454 |
| | 0.0838 | 0.067 | 0.101 | 0.264 | |

Data expressed as mean relative expression ($2^{-\Delta Ct}$) ± SEM. WIS: Wistar, WKY-Veh: Wistar Kyoto-Vehicle, WKY-Sert: Wistar Kyoto-Sertraline, WKY-Ex: Wistar Kyoto-Exercise. P-value derived from One-Way ANOVA.

**Supplementary Table 3: Dam behavioural data from WKY-Ex high runners and low runners**

| | Low Runners | High Runners | T-test |
|---|---|---|---|
| Time Spent in Open Arms (EPM) | 37.67± 11.25 | 21.49± 7.99 | t=-1.172, p=0.285 |
| Time Spent in Closed Arms (EPM) | 215.98± 29.65 | 249.35± 27.96 | t=0.819, p=0.444 |
| Distance Travelled (OFT) | 3095.65± 273.12 | 3067.18± 103.05 | t=-0.098, p=0.925 |
| Time Spent in corners (OFT) | 435.83± 38.44 | 435.30± 25.43 | t=-0.012, p=0.991 |
| Time Spent in Centre (OFT) | 74.20± 15.90 | 105.68± 18.26 | t=1.300, p=0.241 |
| **Novel Object Interaction Time (NOR)** | **0.56± 0.05** | **0.77± 0.01** | **t=4.275, p=0.005** |
| Sucrose Preference | 75.75± 3.33 | 77.09± 5.74 | t=0.202, p=0.846 |

High runners represent those 4 dams who ran more than the median running distance. Low runners represent those 4 dams who ran less than the median running distance. Data presented as Mean ±SEM. EPM: Elevated Plus Maze; NOR: Novel Object Recognition; OFT: Open Field Test.

**Supplementary Table 4: Pup Retrieval Test Data**

| Dam's Group | Pick 1st pup | Return 1st pup to the nest | Return all the pups to the nest |
|---|---|---|---|
| WIS | 4.0/6.0 | 2.0/6.0 | 0.00/6 |
| WKY-Veh | 6.0/8.0 | 6.0/8.0 | 3.0/8.0 |
| WKY-Sert | 4.0/5.0 | 4.0/5.0 | 2.0/5.0 |
| WKY-Ex | 6.0/8.0 | 6.0/8.0 | 6.0/8.0 |

Pup retrieval test at PN7. Number of pups' retrieval task completers vs non-completer of: 1) pick up 1st pup by dam's mouth, 2) return at least one pup to the nest, and 3) return all the litter in their nest, for Wistar (Wis; n=6), Wistar Kyoto Vehicle (WKY-Veh; n=8), Wistar Kyoto sertraline (WKY-Sert; n=5), and Wistar Kyoto exercise (WKY-Ex; n=8) groups.
